# Emergence of V1 connectivity pattern and Hebbian rule in a performance-optimized artificial neural network

**DOI:** 10.1101/244350

**Authors:** Fangzhou Liao, Xiaolin Hu, Sen Song

## Abstract

The connectivity pattern and function of the recurrent connections in the primary visual cortex (V1) have been studied for a long time. But the underlying mechanism remains elusive. We hypothesize that the recurrent connectivity is a result of performance optimization in recognizing images. To test this idea, we added recurrent connections within the first convolutional layer in a standard convolutional neural network, mimicking the recurrent connections in the V1, then trained the network for image classification using the back-propagation algorithm. We found that the trained connectivity pattern was similar to those discovered in biological experiments. According to their connectivity, the neurons were categorized into simple and complex neurons. The recurrent synaptic weight between two simple neurons is determined by the inner product of their receptive fields, which is consistent with the Hebbian rule. Functionally, the recurrent connections linearly amplify the feedforward inputs to simple neurons and determine the properties of complex neurons. The agreement between the model results and biological findings suggests that it is possible to use deep learning to further our understanding of the connectome.

## 1 Introduction

The mono-synaptic excitatory recurrent (or lateral) circuit in layer 2/3 (Gilbert and Wiesel, 1983) and layer 4 (Li et al., 2013) of V1 is one of the most well-studied circuit in the cortex. In some species like ferret (Bosking et al., 1997) and cat (Schmidt et al., 1997), it is found that neurons in different columns with collinearly located receptive fields (RF) tend to have long-range connections. The short-range connections are harder to be studied, and only a few studies were carried out in mice mice (Ko et al., 2011, 2013, Cossell et al., 2015), which does not have the columnar organization in V1. It was found that the neurons with similar orientation tuning and high RF correlation are more likely to be connected (Ko et al., 2011, Lee et al., 2016), and tend to have strong synaptic weight (Cossell et al., 2015).

At the physiological level, based on the anatomical evidences described above, it is widely accepted that the excitatory recurrent connections contribute to collinear facilitation (Nelson and Frost, 1985, Kapadia et al., 1995, Chisum et al., 2003). They are also hypothesized to linearly amplify the feedforward input, which is supported by both theoretical analysis (Martin and Suarezt, 1995) and experimental evidence (Li et al., 2013). The full recurrent circuit (excitatory and inhibitory) also take part in surround suppression (Weliky et al., 1995, Adesnik et al., 2012). At the perceptual level, some studies proposed that they take part in functions like contour integration (Li, 1998) and saliency map (Li and Gilbert, 2002). Adini et al. (2002) used recurrent connections to explain the improved contrast discriminating performance after practicing in the presence of similar laterally placed stimuli. But it is unclear if it is these factors that entail the recurrent connectivity pattern because these studies only show that recurrent connections enable such computations. A computational model that explains the emergence of the recurrent connectivity pattern is still lacked. In this study, we explore this problem by assuming that the recurrent connectivity is a result of performance optimization in visual recognition.

The recent rising of the artificial neural networks gives us an inspiration. The convolutional neural network (CNN) shows many similarities with the biological visual system. Structurally, both of them are composed of hierarchical layers, and each layer is composed of many units (columns), different units process information in different locations. Functionally, most kernels of the first convolutional layer are Gabor filters, resembling the RF of V1 neurons (Krizhevsky et al., 2012), the feature representation in higher-level is correlated with that in area V4 and IT (inferior temporal) cortex (Yamins et al., 2014). In *k*-categories image recognition task with N training samples, the synaptic weights of CNN, denoted by *w*, are usually obtained by minimizing the cost function:

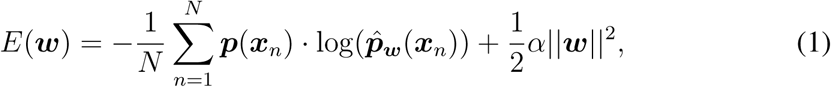
where *p*(*x_n_*) is a *k*-dimensional one-hot vector and stands for the ground truth provided for the sample *x_n_*, and 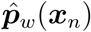 is the predicted answer given by the CNN. The first term measures on average how close the answer is to the ground truth. The second term encourages small weights by using the *l*_2_ norm regularization, which imposes weight decay to all connection weights during training. It is a standard method for preventing overfitting. In other words, the goal of CNN is two-folded: “better fitting” and “simpler structure”. The coefficient *α* controls the weights of these two goals.

We hypothesize that during millions of years of evolution, the brain also pursues these two goals. Thus the functional similarity between the brain and CNN may root in the similarity between their cost functions. Furthermore, if we introduce recurrent connections in the CNN and optimize it for image classification, its connectivity might also be similar to that of the brain. In this work, we adopted a CNN incorporated with recurrent connections in the first layer, and trained it for image classification task. It was found that both the resultant connectivity and the functional roles of the recurrent connections in the first layer were very similar to that of V1.

### 1.1 Related work

Neuroscience and deep learning can benefit each other (see Hassabis et al. (2017) for a recent review). In particular, in recent years, deep neural networks have been used to reveal functional mechanisms of the brain. Yamins et al. (2014) showed that the responses of V4 and IT neurons can be linearly regressed with the responses of a performance-optimized deep network. Khaligh-Razavi and Kriegeskorte (2014) compared tens of computational models and found that supervised deep models better explain IT cortical representation for natural images than unsupervised models. Based on experiments on pretrained deep neural networks on large image datasets, Zhuang et al. (2017) predicted the correlation of neural response sparseness and a functional signature of higher areas in the visual pathway. These studies focus on the functional properties of neurons and did not study the connectivity patterns among neurons.

Song et al. (2016) investigated the connectivity of a recurrent network that is optimized in several cognitive tasks. But first, although they got a full connection matrix, they did not find a clear connectivity pattern in the neuronal level; second, there is no certain corresponding brain region for their model, so it is hard to confirm their connectivity predictions in biological experiments.

Liang and Hu (2015) proposed the recurrent convolutional neural network (RCNN) and showed that the extra recurrent connections can improve the classification accuracy of the standard CNN. Spoerer et al. (2017) showed that RCNN performed significantly better in object recognition task under occlusion. Liao and Poggio (2016) demonstrated the relationship between recurrent circuits and the Residual network (He et al., 2016). McIntosh et al. (2016) built a CNN to model retina response data, and found that the introduction of lateral connection in their model enabled it to capture contrast adaptation. But the recurrent connectivity pattern in all of these works was not discussed.

## 2 Methods

### 2.1 Dataset

For the ease of training, we selected 50 classes of images from the Imagenet dataset (Deng et al., 2009), resulting in a dataset with 64494 training samples and 2500 validation samples. In addition, we resized images to 64 × 64 pixels and converted them to grayscale because we did not intend to analyze the color information.

### 2.2 Network Design

First, we designed a feedforward network as the baseline. Its first convolutional layer has 64 7 × 7 kernels, and other details of the architecture are shown in Table 1. To model the recurrent connection, we adopted a modified version of RCNN (Liang and Hu, 2015). An extra 7 × 7 convolutional layer which receives input from the first layer and then sends output back is designed to model the recurrent connections in the first layer. Since the number of parameter of this recurrent convolution kernel is too large (7 × 7 × 64 × 64) to be fully optimized, it is factorized as a channel-wise 7 × 7 convolution and a cross-channel 1 × 1 convolution. The number of parameters of the recurrent layer is reduced to 7 × 7 × 64 + 64 × 64 via this simplification. The 1 × 1 convolution simulates intra-column connection, and the channel-wise convolution simulates the inter-column connection (Figure 1a). The simplification means that a neuron does not connect to another neuron with a different RF shape at a different location. According to biological finding (Gilbert and Wiesel, 1989), this simplification is reasonable. The network architecture is shown in Figure 1b.

**Figure 1:**
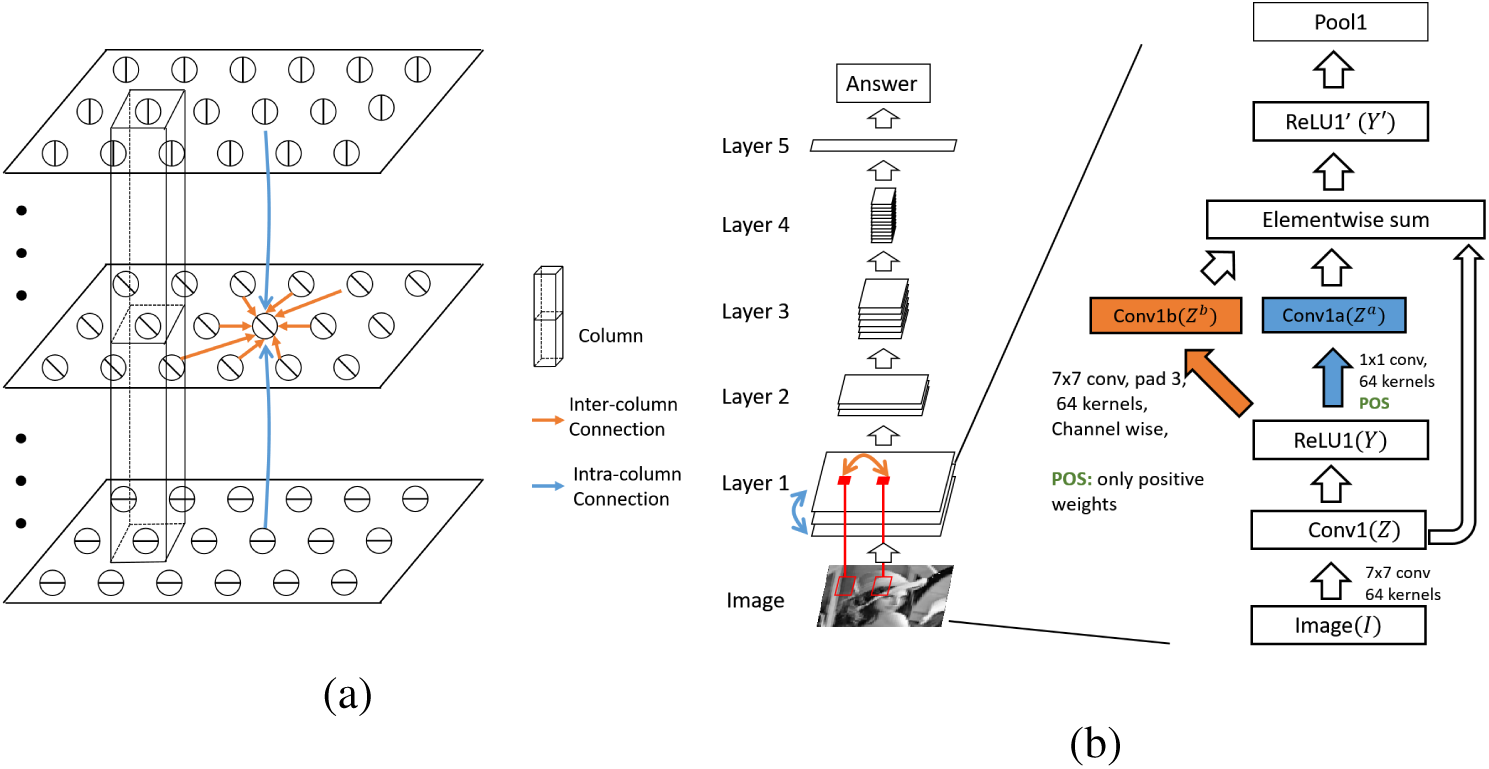
The architecture of the model. (a) The diagram of the first convolutional layer. Three large rectangles stand for three different feature maps. Note the columnar structure and intra/inter-column connections. (b) Left panel: The structure of the whole network which has 5 layers. Recurrent connections are introduced in the first layer. Right panel: The designing details of the first layer. Conv: convolution; ReLU: rectified linear unit; Pool: pooling.

The computation process in the first layer can be written as:

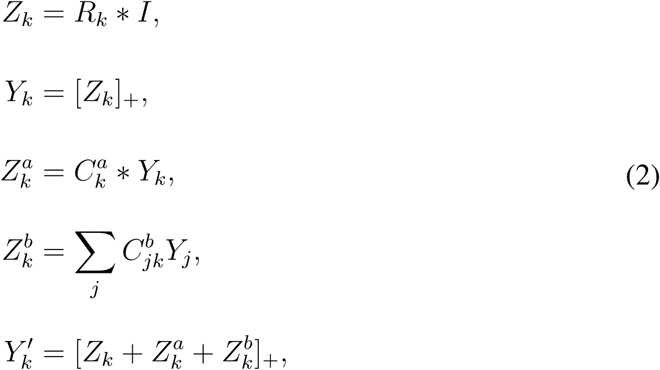
where * denotes the convolution operation, *k* is the index of the feature map, [·]_+_ denotes the rectify linear function, *R* ∈ **R**^64×7×7^ denotes the feedforward convolutional kernels (also called feedforward RFs), *C^a^* ∈ **R**^64×7×7^ denotes the inter-column connection, *C^b^* ∈ **R**^64×64^ denotes the intra-column connection.

For convenience, we name the feedforward model as FFM and the model with recurrent connections as RCM.

### 2.3 Optimization & Constraints

The gradients of parameters with respect to the loss function (1) are calculated with back-propagation algorithm (Rumelhart et al., 1986), and the stochastic gradient descent method was used to update the parameters. The initial learning rate was set to 0.01 for the first 100 epochs and decayed by a factor 0.1 every 100 epochs. A total of 400 epochs was used. The momentum was set to 0.9, and the coefficient *α* was set to 0.0005.

**Table 1:** The architecture of the baseline model

| Layer | Data | Conv1 | Pool1 | Conv2 | Pool2 | Conv3 | Pool3 | Conv4 | Pool4 | Conv5 | Pool5 | Classifier |
| --- | --- | --- | --- | --- | --- | --- | --- | --- | --- | --- | --- | --- |
| Param | Mean | 7x7, Pad 3 | Max, 2x2 | 3x3, Pad 1 | Max, 2x2 | 3x3, Pad 1 | Max, 2x2 | 3x3, Pad 1 | Max, 2x2 | 3x3, Pad 0 | Ave, 2x2 | Softmax |
|  | subtraction | St 1, ReLU | St 2 | St 1, ReLU | St 2 | St 1, ReLU | St 2 | St 1, ReLU | St 2 | St 1, ReLU | St 2, Drop 0.5 |  |
| Size | 64x64x1 | 64x64x64 | 32x32x64 | 32x32x64 | 16x16x64 | 16x16x128 | 8x8x128 | 4x4x256 | 4x4x256 | 2x2x384 | 1x1x384 | 1x1x50 |

St: stride, Drop: dropout ratio, Ave/Max: average/max pooling, Pad: padding

Our goal is to simulate the function of pyramid neurons and the connections between them, so the recurrent connections (*C^a^*, *C^b^*) were constrained to be positive. In addition, the self-connections 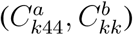 were set to zero. These constraints were achieved by setting all negative weights and self-connections to zero after every updating iteration.

## 3 Results

### 3.1 Feedforward RFs

The RFs of all neurons in the first layer in the model are shown in Figure 2. There were 13 neurons with extremely low RF amplitudes and irregular RF shapes. They are called “complex” cells, and the reason will be discussed later. The others are called “simple” neurons.

**Figure 2:**
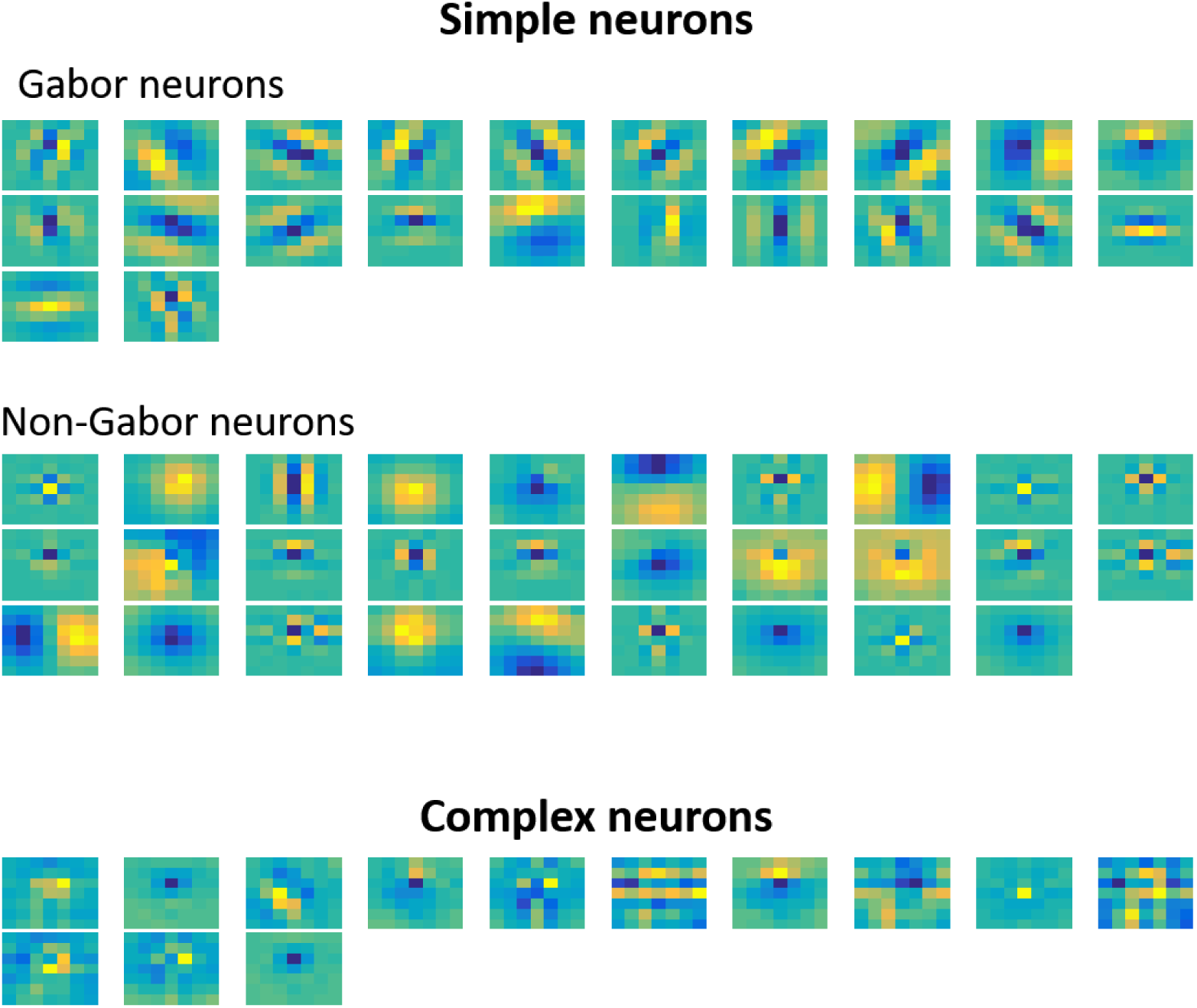
The feedforward RFs of all neurons.

We fitted the RFs of all simple neurons with the Gabor function. We found that some 7 × 7 kernels were over-fitted, so we padded all kernels to 11 × 11 with 0 before fitting. Based on the fitting error, orientation selectivity, spatial frequency limitation, we selected 22 Gabor-like RFs, and named them Gabor neurons. The other 29 neurons are called Non-Gabor neurons.

### 3.2 Connectivity among layer one neurons

#### 3.1 Intra-column connections

We first extracted the preferred orientation of every Gabor neuron. It was found that neurons with closer preferred orientation tend to have larger synaptic weights (Figure 3a). Then we fitted the synaptic weights with RF correlation. Consistent with the biological findings by Cossell et al. (2015), the latter fitting was better than the former (Figure 3b). An even better fitting was obtained using the inner product of the two RFs as it explained most variance (*r*^2^ = 0.86, Figure 3c). The fitting equation is

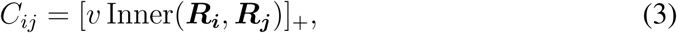
where ***R****_i_* denotes the RF of the *i*-th neuron, *C_ij_* denotes the synaptic weight from *j*-th neuron to *i*-th neuron, Inner(*x, y*) stands for the inner product between × and *y*, and [ ]_+_ stands for the rectifier function max(0, *x*). Because the mean value of ***R****_i_* was close to 0, the equation can be equivalently written as:

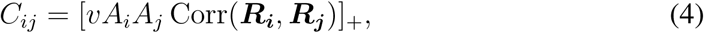
where Corr(*x, y*) denotes the correlation between *x* and *y*, and *A_i_* denotes the amplitude or the norm of ***R****_i_*. It means that the neurons with higher average firing rate and more similar receptive fields tend to have larger synaptic weight, which exactly reflects the Hebbian rule.

**Figure 3:**
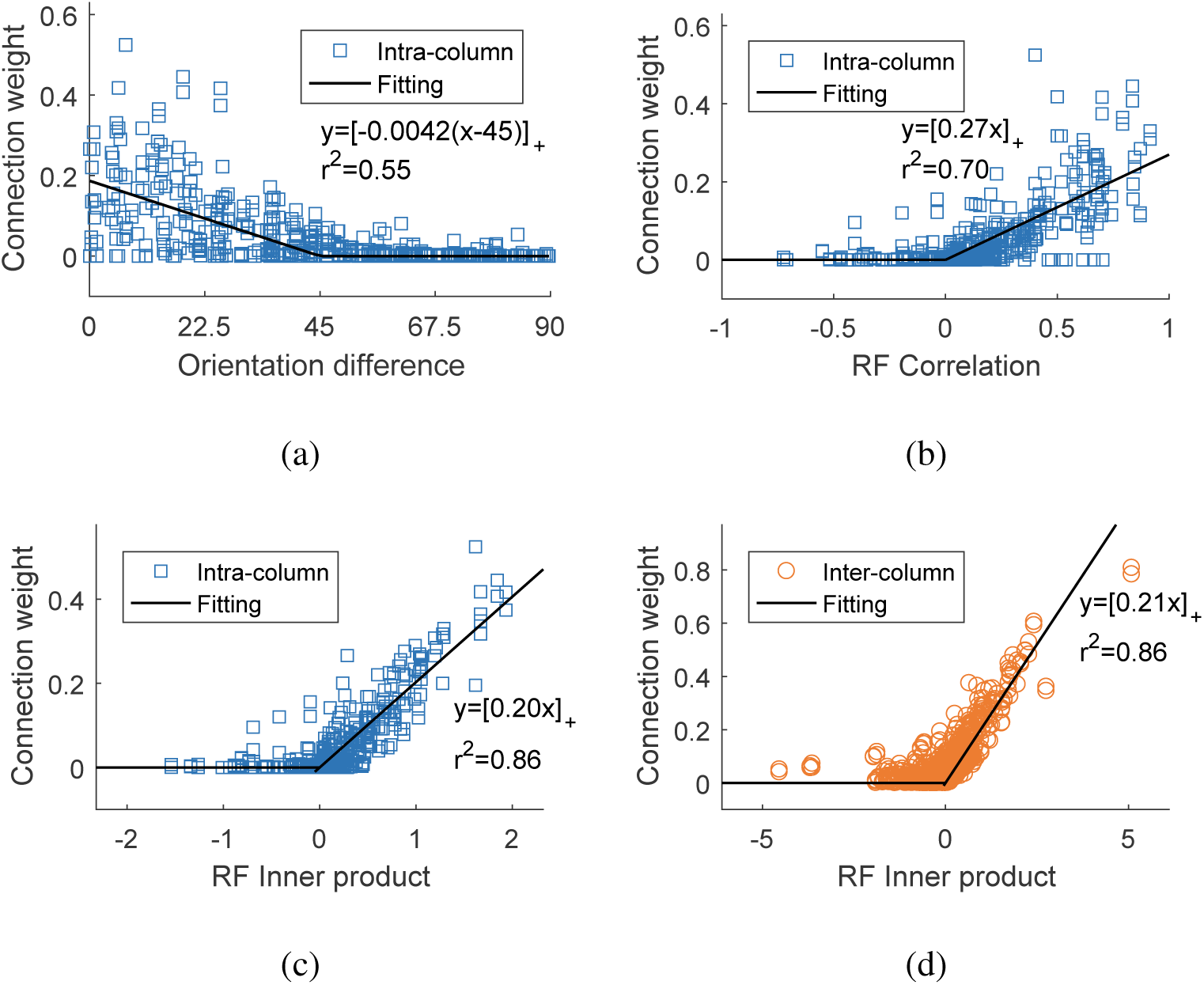
The statistical connectivity pattern of the Gabor neurons in the model. (a-c) Fitting intra-column connection weights of Gabor neurons with preferred orientation difference (a), RF correlation (b), and RF inner product (c). (d) The same as (c) for inter-column connections of the Gabor neurons.

#### 3.2 Inter-column connections

Notice that because we factorized the convolution operation, each neuron only receives inter-column signals from other neurons with the same RF shape. So each feedforward convolution kernel connecting the input image and a feature map corresponds to an inter-column convolution kernel in that feature map. The corresponding feedforward RFs and inter-column recurrent kernels are shown in the left panel of Figure 4a. Notice that the orientations of recurrent connection kernels are very similar to those of their corresponding feedforward convolution kernels, indicating that the neurons with collinear RFs tend to excite each other, a phenomenon called collinear facilitation for V1 neurons (Bosking et al., 1997). See the right panel of Figure 4a for an illustration.

**Figure 4:**
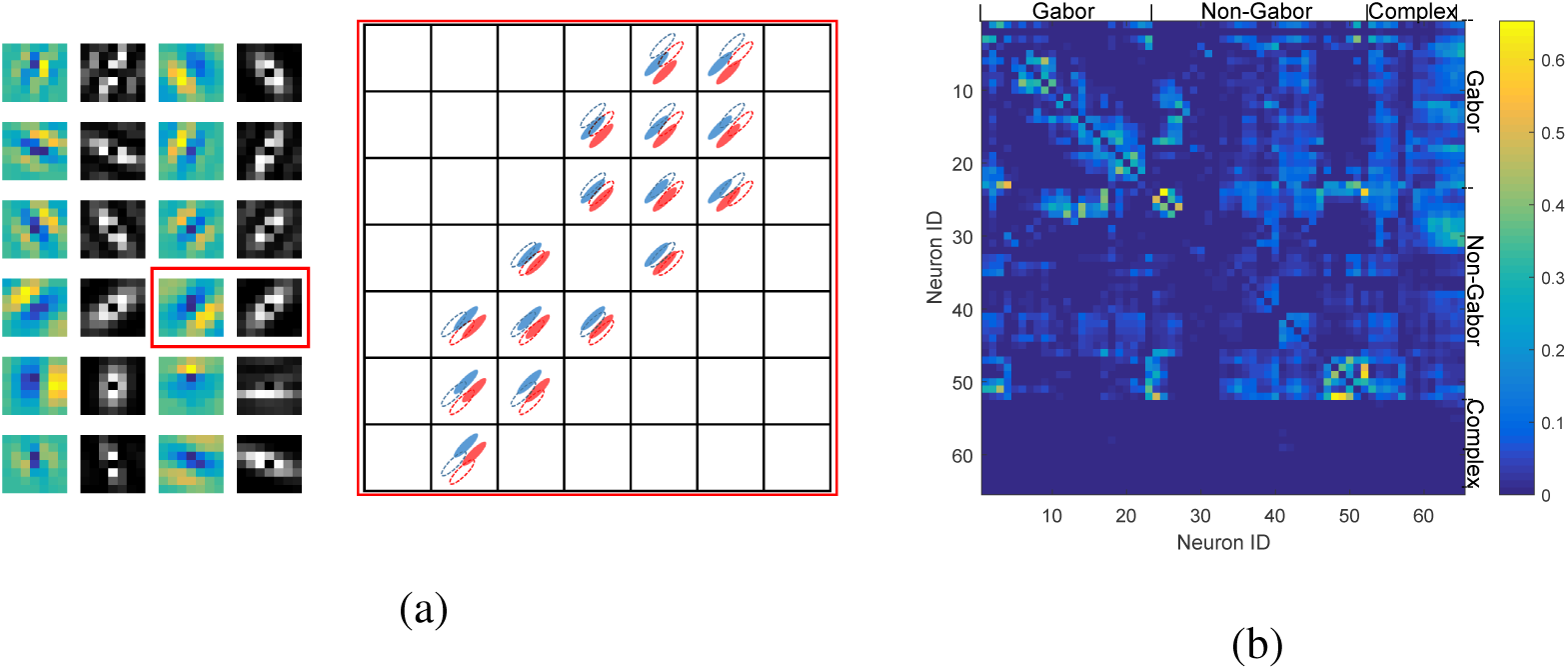
The illustration of the recurrent connection pattern. (a) The illustration of the inter-column connection. Left panel: the colorful images show the RF of Gabor neurons, and the gray scale images show the inter-column connection matrix of its corresponding RF. Right panel: Enlarged illustration for the inter-column connection matrix highlighted in the left panel. Each grid block corresponds to an entry of recurrent connection matrix and stands for the connection strength between a neuron pair. The distance between a grid block and the central grid block stands for the displacement between the RFs of pre- and post-synaptic neurons. In each grid, the dashed line RF and filled RF stand for the pre- and post-synaptic neurons, respectively. Only the strong connections (stronger than 0.2 times the strongest connection) are shown. (b) The intra-column connection matrix of all neurons. Each row stands for the output connections of a neuron, and each column stands for the input connections of a neuron. The Gabor neurons are sorted by their preferred orientations, while non-Gabor and complex neurons are clustered based on their connectivity similarity.

We then investigated the quantitative relationship between the inter-column connection weights and the corresponding pairs of RFs. Note that the RF shapes of all neurons in the same feature map are the same but their locations are different. So to make a pair of RFs match each other in spatial location, they are shifted and padded with zeros. Using these RFs, it was found that the inter-column connections also satisfied Equation (4) (Figure 3d). Even the fitting coefficients are close to that of intra-column connections.

#### 3.3 Emergence of “complex” neurons

The full connection matrix of intra-column connections is shown in Figure 4b. Most neurons had quite symmetric intra-column synaptic weights. But some neurons only received intra-column recurrent inputs but did not send output. In addition, the amplitudes of their feedforward RFs were very low, so the feedforward activation was almost zero. In other words, the properties of this kind of neuron are solely determined by other neurons connected to it by intra-column connections. This connection pattern is similar to that of complex neurons in V1, so we call them “complex” neurons ^1^.

For other non-Gabor neurons, most of them had similar properties with Gabor neurons: symmetric connections, clustering with similar neurons, and satisfying Equation 3. in other words, they are simple cells without perfect Gabor RFs.

#### 3.4 Non-random connectivity pattern

Song et al. (2005) found that one notable feature of the recurrent connections in the cortex is their non-random motifs. That is to say, the frequency of some two-neuron and three-neuron connectivity patterns (motifs) is higher than that of a randomly connected network. We conducted the same analysis in our model using the whole intra-column connection matrix.

First, to model the binary connection in the original study (Song et al., 2005), we binarized the connection matrix. The basic idea of binarizing is that the connections with larger weights are more likely to be kept:

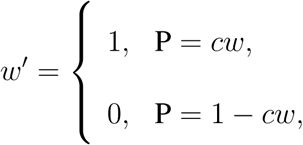
where *w* is the synaptic weight, *w*′ is the binarized weight, and *c* is a probability normalization factor. *c* was adjusted so that the overall connection probability was 11.6%, which is consistent with the original research (Song et al., 2005). The distribution of the initial weights (*w*) and chosen weights (*w′w*) are shown in Figure 5a. Then we counted the frequency of all two-neuron and three-neuron patterns in this network relative to those in a random network. The ratio between the two frequencies for each motif reflects “non-randomness” of the motif in our network. The results of the two-neuron motifs were very similar to physiological data, and the results of several three-neuron motifs (such as #14,15,16), but not all (such as #5,11,12), were similar to physiological data.

**Figure 5:**
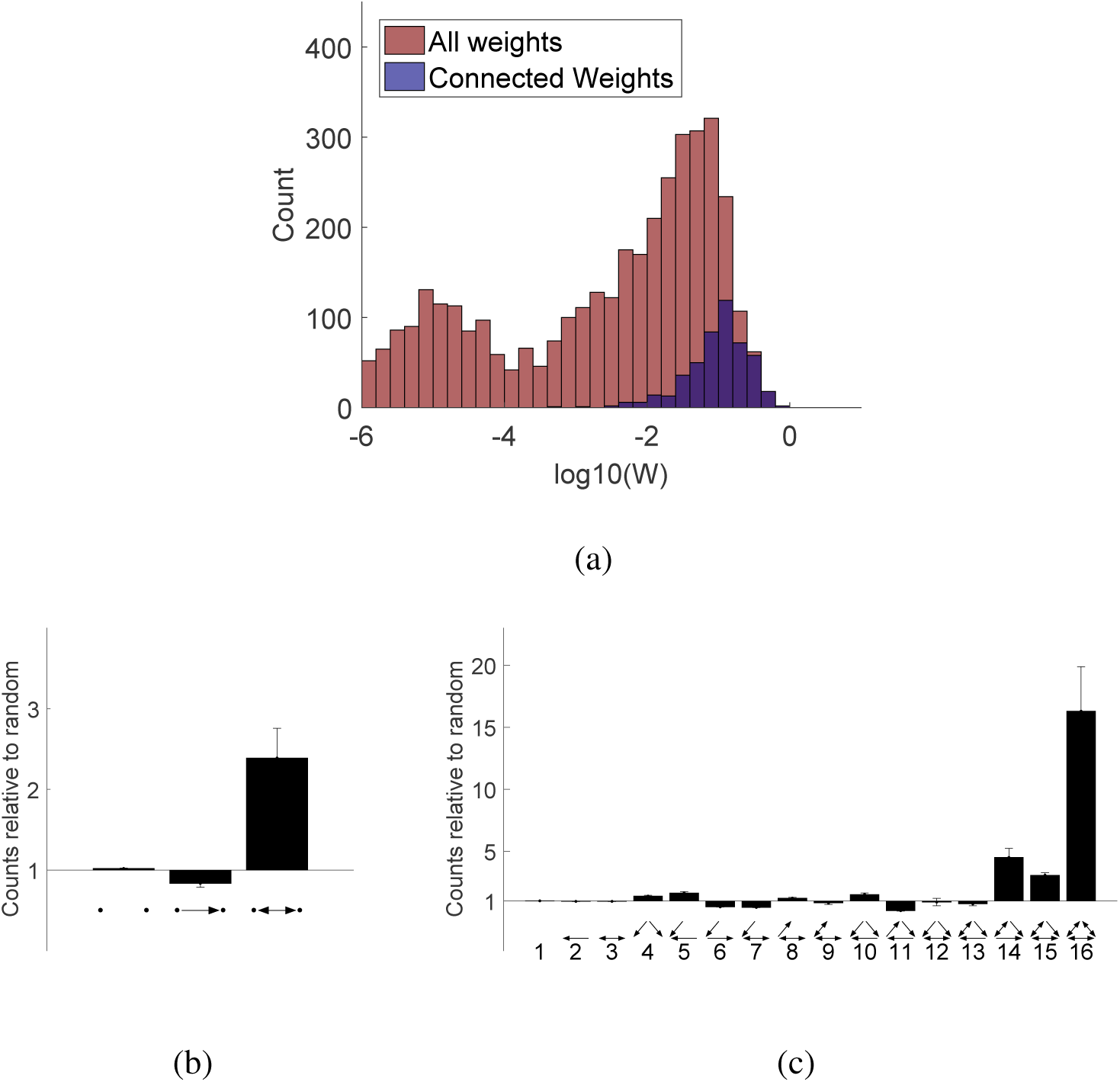
Emergence of nonrandom connectivity patterns. (a) The distribution of the weights in intra-column recurrent connection matrix and the weights chosen to be connected for motif analysis. The *x*-axis is in log-scale. (b) The reciprocal connection probability is higher than random probability. (c) The three-neuron patterns probability relative to the random network. The results in (b) and (c) are averaged from four independent experiments, the error bars indicate the standard deviation.

### 3.3 Functional roles of recurrent connections

To investigate the effect of recurrent computation, we compared the activation of a typical Gabor neuron before and after recurrent computation (Figure 6a). It was found that the total activation was basically a linear amplification of the feedforward activation for Gabor neurons except for one outlier, and the amplifying ratio was around 3.5 (Figure 6b). This phenomenon was also observed in the layer 4 neurons in V1 (Li et al., 2013).

**Figure 6:**
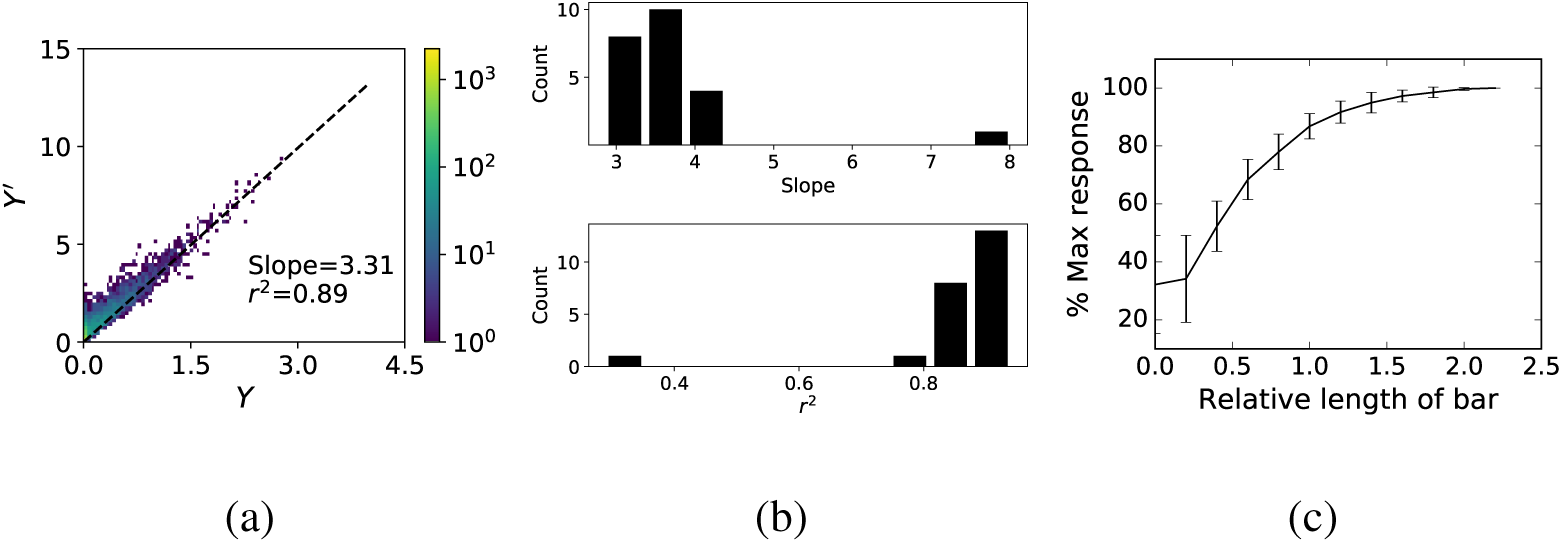
Functional roles of the recurrent connections. (a) The activity of a typical Gabor neuron before (*Y*) and after (*Y*′) the recurrent computation. The color stands for data density. (b) The summary of results of linearly fitting the activation before and after recurrent computation for all Gabor neurons. Upper panel: the histogram of amplifying ratio (slope). Lower panel: the histogram of the correlation coefficient. The outlier in the upper panel corresponds to that in the lower panel. (c) The collinear facilitation effect. The *x*-axis indicates the bar length relative to the length of the RF. The *y*-axis indicates the response relative to the response to the longest bar. The data is averaged across all Gabor neurons, the error bars indicate the standard deviation.

Another well-known function of recurrent connections is collinear facilitation. We examined the response of each Gabor neuron when it was presented with a bar stimulus with the neuron’s preferred orientation and location. The relationship between the activity and the bar length relative to the RF length are plotted in (Figure 6c). The activation of the Gabor neuron after recurrent computation still grows when the length of the bar goes beyond the RF length. The length of summation field was about 2 times as the RF length, and the response magnitude could be improved by around 20% by collinear facilitation.

The magnitude of collinear facilitation in our model was slightly weaker than that found in experiments (Chisum et al., 2003, Angelucci and Bressloff, 2006). The reason for this discrepancy might be that RF mapped in the biological experiments tend to be smaller than true RF due to high noise in biological experiments.

To visualize the collinear facilitation directly, we feed a dashed line image to the network, and choose a neuron whose preferred orientation is the same as the dashed line. The representation of the chosen neuron (its feature map) is shown in Figure 7. Before recurrent computation, the neurons whose RF located between two dashes has nearly 0 activation. But after recurrent computation, the activations of these neurons are strengthened so that the representation of the whole dashed line becomes more continuous. This effect is called contour completion.

**Figure 7:**
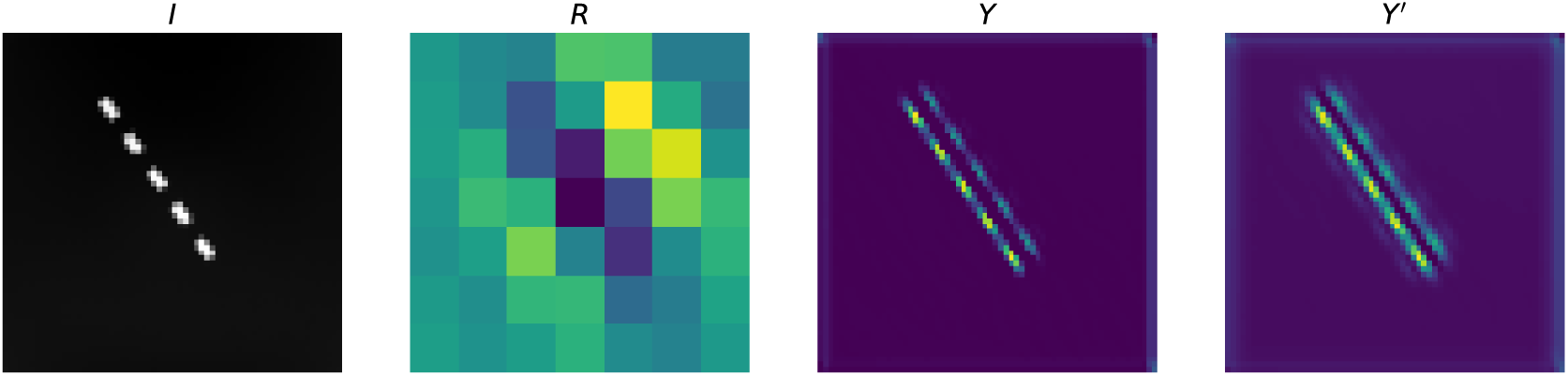
Contour completion effect. Left to right: the input image; the RF of the inspecting neuron; the representation (feature map) of the inspecting neuron before recurrent computation; the same as the last one, but after recurrent computation.

#### 3.1 Effect of regularization level

We tried different *α* levels (*α* = 0, *α* = 0.0005, *α* = 0.005) in both FFM and RCM, and found that the performance and resultant connectivity strongly depended on *α*. When *α* = 0, FFM showed better performance than RCM. In addition, the relationship between RF inner product and connection weights becomes noisy (Figure 8b). The functional role of recurrent connections was no longer linear amplifying, either. These results suggest that the *l*_2_ regularization term is necessary for the emergence of the recurrent connectivity pattern described above. When *α* = 0.0005, the RCM slightly outperformed FFM. When *α* = 0.005, RCM performed significantly better than FFM (Figure 8a). That is to say, recurrent connection plays more important role in models with higher *α*. We provide a more detailed explanation in the next section.

**Figure 8:**
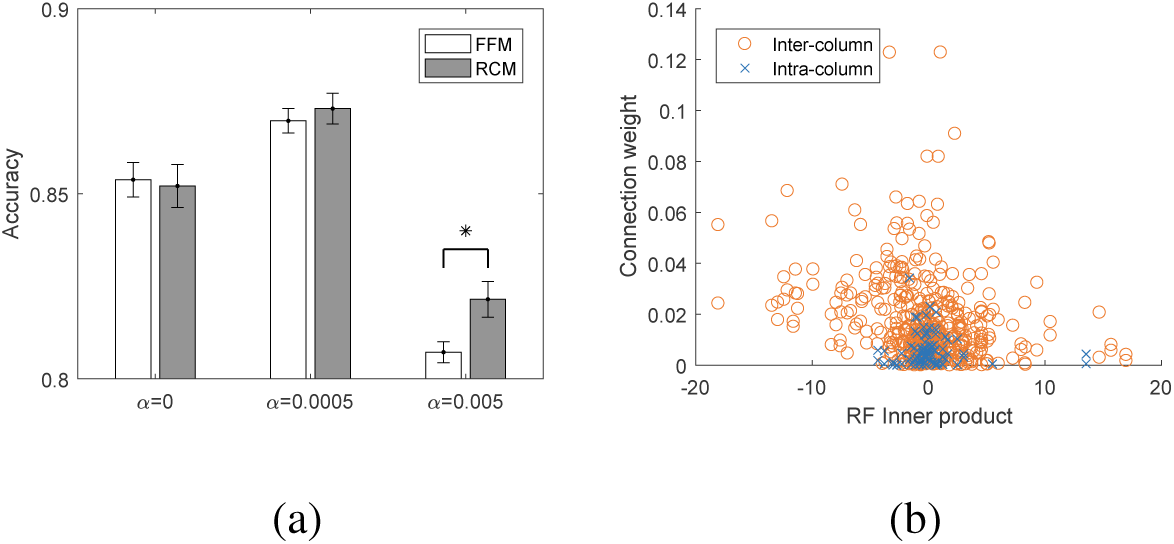
The influence of the regularization factor. (a) The recognition performance of FFM and RCM under different regularization level (*α*). The error bars indicate the standard deviation. N=4 for each condition. (b) The recurrent connection pattern found in RCM without the regularization term. The analysis is the same as that for Figure 3c and 3d.

## 4 Discussion

In this work, we optimized a CNN equipped with monosynaptic excitatory recurrent connections by purely supervised learning and found that the recurrent connection pattern is similar to which discovered in the brain. And the resultant synaptic weights can be explained by Hebbian rule, which improves the biological plausibility of our results.

### 4.1 The existence of *A_i_A_j_*

Cossell et al. (2015) have revealed that synaptic weight between a pair of V1 neurons is related to the correlation of their RFs. We introduce new factors: the amplitudes of RFs, *A_i_* and *A_j_* in Equation (4). So a natural question is whether these two factors are reasonable. Due to technical difficulties, this hypothesis can not be tested directly by experiment. Fortunately, we still have two pieces of indirect supporting evidence in biological experiments. First, the fitting in the original report (Cossell et al., 2015) was not good enough—the weights between some neuron pairs with high RF correlation were very low. This can be explained by introducing a low amplitude to one of these neurons in our model. Second, if V1 neurons are categorized into “responsive” and “unresponsive” neurons according to their responsiveness to visual stimuli, the connection probability among the “responsive” neurons is found to be higher than that among “unresponsive” neurons (Mrsic-Flogel, 2012). This is consistent with Equation (4) because *A_i_* and *A_j_* reflect the responsiveness.

### 4.2 Recurrent connections can reduce *l*_2_ cost

In the FFM, we added an *l*_2_ regularization term to the loss function. The parameter *α* controls the trade-off between fitting variance and bias. A proper *α* value can reduce the fitting variance and thus improve generalization ability, but when *α* grows too big, the fitting is biased and the weights after training are discounted (left panel of Figure 9).

**Figure 9:**
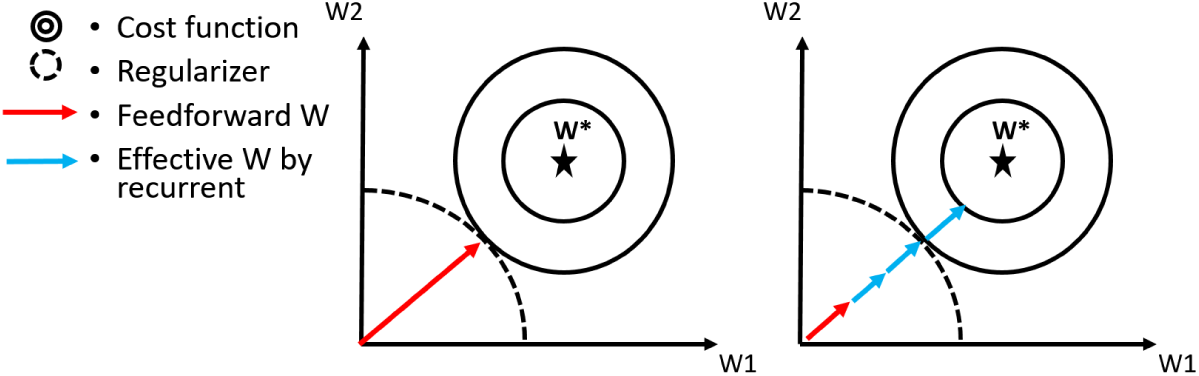
An illustration of the effect of recurrent connections. Left panel: in FFM, the presence of large *l*_2_ regularization makes the trained parameter strongly discounted. Right panel: in RCM, the recurrent connections act as a linear amplifier of the feedforward connections, and the weights can be less discounted with the same total *l_2_* cost as in the FFM.

In the RCM, a layer 1 neuron connects to neurons with similar RFs. In effect, these connections act as extra feedforward paths, compensate the weight decay effect and remedy the fitting bias (right panel of Figure 9). The recurrent connections also suffer from weight decay, but the total *l_2_* cost of RCM can be smaller than that of FFM. Because the *l*_2_ term punishes the large synapses more, breaking a large synapse into several smaller synapse can be economical. To see this, consider a feedforward system composed of two neurons with the same RF *w*. An equivalent recurrent system is composed of two neurons with 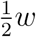 and reciprocal connection 1. The total *l*_2_ terms for these two systems are 2||*w*||^2^ and 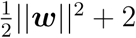 respectively. So when ||*w*||^2^ is large enough, the recurrent connections are economical. This effect is even more significant in a large system, and that is why the amplification ratio in our model is so high (Figure 6b). There is another factor that favors recurrent connections: in the brain, feedforward connections are much longer than recurrent connections, thus the maintaining and transferring cost on feedforward connections may be larger than that on recurrent connections.

### 4.3 Limitations

As a preliminary exploration, this study has many limitations. First, only supervised learning is considered but it is well-known that unsupervised learning is also very important for the survival of animals. We could only argue that supervised learning can lead to the experimentally found recurrent connectivity pattern in V1, but do not exclude the contribution of unsupervised learning in shaping this pattern.

Second, biological plausibility of the back-propagation algorithm used in training the model is controversial. But the point we want to emphasize in this study is the loss function, not the specific optimizing algorithm. Biological system may use another algorithm to achieve the same solution.

Finally, for technical reasons (training a deep model on a large number of images as in this study is not an easy task), we have made some simplifications in modeling. For example, inhibitory neurons were not incorporated in the model, recurrent connections were factorized into inter-column and intra-column part, and the number of recurrent steps was only one. So some important roles of recurrent connections, such as surround suppression and normalization, can not be explained by this model.

## Acknowledgements

We thank Zhe Li for useful discussion. This work was supported in part by the National Natural Science Foundation of China under grant nos. 61332007, 61621136008 and 61620106010.

## Footnotes

1 Usually the max-pooling neuron is considered to act as complex neuron because they have translation invariance. Not all of the “complex” neurons in this study have this property. We use the term “complex” just because these neurons cannot be described by simple linear filter.

